# In silico biophysics and rheology of blood and red blood cells in Gaucher Disease

**DOI:** 10.1101/2024.12.10.627687

**Authors:** Zhaojie Chai, Guansheng Li, Papa Alioune Ndour, Philippe Connes, Pierre A. Buffet, Melanie Franco, George Em Karniadakis

## Abstract

Gaucher Disease (GD) is a rare genetic disorder characterized by a deficiency in the enzyme glucocerebrosidase, leading to the accumulation of glucosylceramide in various cells, including red blood cells (RBCs). This accumulation results in altered biomechanical properties and rheological behavior of RBCs, which may play an important role in blood rheology and the development of bone infarcts, avascular necrosis (AVN) and other bone diseases associated with GD. In this study, dissipative particle dynamics (DPD) simulations are employed to investigate the biomechanics and rheology of blood and RBCs in GD under various flow conditions. The model incorporates the unique characteristics of GD RBCs, such as decreased deformability and increased aggregation properties, and aims to capture the resulting changes in RBC biophysics and blood viscosity. This study is the first to explore the Young’s modulus and aggregation parameters of GD RBCs by validating simulations with confocal imaging and experimental RBC disaggregation thresholds. Through *in silico* simulations, we examine the impact of hematocrit, RBC disaggregation threshold, and cell stiffness on blood viscosity in GD. The results reveal three distinct domains of GD blood viscosity based on shear rate: the aggregation domain, where the RBC disaggregation threshold predominantly influences blood viscosity; the transition area, where both RBC aggregation and stiffness impact on blood viscosity; and the stiffness domain, where the stiffness of RBCs emerges as the primary determinant of blood viscosity. By quantitatively assessing RBC deformability, RBC disaggregation threshold, and blood viscosity in relation to bone disease, we find that the RBC aggregation properties, as well as their deformability and blood viscosity, may contribute to its onset. These findings enhance our understanding of how changes in RBC properties impact on blood viscosity and may affect bone health, offering a partial explanation for the bone complications observed in GD patients.

**Author summary:** In Gaucher Disease (GD), a genetic deficiency in the enzyme glucocerebrosidase leads to the accumulation of glucosylceramide in red blood cells (RBCs), resulting in altered biomechanical properties. These changes affect blood flow characteristics, particularly blood viscosity, and may contribute to bone health issues seen in GD patients, including bone infarcts, avascular necrosis (AVN), and other bone diseases. In our study, we apply dissipative particle dynamics (DPD) simulations to explore how GD impacts RBC behavior under various flow conditions. We model GD RBCs with decreased deformability and increased aggregation, examining how these properties influence blood viscosity across three distinct shear rate domains: aggregation, transition, and stiffness. By validating our simulations with confocal imaging data and experimental RBC disaggregation thresholds, we quantitatively assess the effects of RBC stiffness, aggregation, and hematocrit levels on blood flow in GD. We find that the RBC aggregation properties, deformability and blood viscosity, may contribute to the onset of bone disease. These findings improve our understanding of how changes in RBC properties influence blood viscosity and may contribute to bone health issues, providing a partial explanation for the bone complications observed in GD patients.

## Introduction

Gaucher Disease (GD) is a rare genetic disorder resulting from a deficiency in glucocerebrosidase (GCase) and stands as the most common autosomal recessive lysosomal storage disorder. The enzyme deficiency is caused by a biallelic mutation in the GBA1 gene [1]. Three types of GD are known. Type 1, which is the most common (90 % of cases in Europe and the USA, but less in other regions), is characterized by hepato-splenomegaly, anemia, thrombocytopenia, complex bone disorders such as osteonecrosis with bone micro-infarcts in ischemic regions, and avascular necrosis (AVN) [2, 3]. The extremely rare Type 2 (acute neuronopathic) and Type 3 (sub-acute neuronopathic) GD, in addition to the symptoms present in Type 1, are associated with severe neurological disorders in Type 2 and variable neurological symptoms in Type [4]. GCase deficiency leads to the accumulation of its substrates, glucosylceramide and secondary sphingolipids, in macrophages. These cells, as natural phagocytes, are responsible for breaking down membrane glycolipids from red blood cells (RBCs) and leukocytes. The Lipid-laden causes macrophages to transform into Gaucher cells (GC) [5]. These GCs have a distinctive phenotype, characterized by a striated “wrinkled tissue paper” appearance [6]. In GD, these chronically, alternatively activated macrophages infiltrate various organs, including the liver, spleen, bone marrow, and sometimes the lungs [1]. The alterations of the monocytic lineage and the rise of GC are considered primary contributors to the major signs of GD [7]. However, several manifestations, such as hepatosplenomegaly, anemia, and bone disorders, may result, at least in part, from the presence of altered RBCs. RBCs from GD patients often exhibit abnormal morphologies, sphingolipid accumulation, reduced deformability, and increased aggregability [8–16]. RBC progenitors display differentiation defects leading to anemia [15].

In GD, RBCs show reduced deformability and higher aggregation properties [8]. These abnormal properties are attributed to the accumulation of glucosylceramide in RBCs [17]. However, investigations into biomechanical properties of RBCs from GD patients and their consequences on blood circulation and bone pathology remain limited [8, 14, 16, 18, 19].

An increase in hematocrit and in RBC membrane’s shear modulus is known to cause a significant increase in blood viscosity [19]. Four main biophysical factors therefore may affect blood viscosity in GD: 1) hematocrit level, 2) RBC deformability, 3) plasma viscosity, and 4) sphingolipid concentration, which may result in attractive interactions between RBCs [8, 20–22]. Hematocrit is higher in asplenic Gaucher patients than in healthy controls and is therefore a potentially important determinant of blood viscosity in GD [20]. By contrast, plasma fibrinogen concentration is similar in GD patients and healthy controls, indicating that increased RBC aggregation likely results from alterations in the RBC membrane than from plasma factors [8]. Plasma viscosity can also be excluded because relevant protein levels in GD are normal [8].

Because experimental *in vitro* measurements rely on rare samples with limited power to validate correlations, we decided to use computational methods to limit experimental variations during sample preparation [23]. Computational methods for studying blood dynamics and rheology include continuum-level models, which describe blood flow using constitutive laws, and discrete particle methods, which reconstruct different blood components. Continuum-level models often use either Newtonian or non-Newtonian representations of blood viscosity but rarely provide systematic sensitivity analyses of biophysical parameters [24]. In contrast, discrete particle methods offer a more accurate model of RBC membranes as coarse-grained particulate material, allowing for precise modeling of cell-cell interactions and accurate recovery of whole blood viscosity [25, 26].

We then investigated GD RBC properties using dissipative particle dynamics (DPD) simulations. DPD accurately modeled blood viscosity by representing RBC membranes as coarse-grained particles [8, 26–31]. This is the first time the DPD method was introduced to study the biophysics and rheology of blood and RBCs in GD. We validated the RBC disaggregation threshold and stiffness with experimental data. The computational model incorporated hemorheological parameters to simulate the behavior of RBCs under different flow conditions. By quantitatively assessing RBC deformability, aggregation, and flow characteristics, we examined the relationship between these parameters and GD pathophysiology, particularly its association with bone infarcts and other bone diseases.

## Materials and methods

### GD RBC model

To investigate the role of GD RBC biophysical factors on blood viscosity, we needed to model the individual RBCs accurately, without compromising the essential hydrodynamics of the system and the convective transport processes that govern blood flow. RBC is a highly deformable biconcave nucleus-free membrane cell with a diameter of ∼8 *µ*m. Known as a viscoelastic object, RBC shows both liquid-like (viscous) and solid-like (elastic) responses to an applied deformation at the same time. This allows RBCs to be very deformable to squeeze through capillaries as small as 3 *µ*m, preserve their shapes at small deformation rates, and orient in the flow direction at larger deformation rates in wider arteries [32–34]. The membrane of RBC is modeled as a set of *N*_*ν*_ DPD particles with the three-dimensional coordinates *X*_*i*_ (*i* ∈ 1, …, *N*_ν_) in a triangulated network of springs with a dashpot on the surface. To model the incompressibility of the RBC membrane, area and volume constraints are applied. Also, considering the bending resistance between all neighboring triangles, the bending rigidity of the membrane can be mimicked.

In this work, we considered the aggregation of normal and GD RBCs under different disaggregation thresholds and stiffness, with the shear modulus and disaggregation threshold of RBCs under these conditions summarized in Table 1. The shear modulus and bending modulus of normal RBCs in our work were selected to be *Es*_0_ = 4.792*µN/m, Eb*_0_ = 2.9 ×10^−19^*J* [35]. We further validated our comprehensive RBC model by comparing it with experimental data, ensuring accurate simulations that capture the changes in biomechanical, rheological, and dynamic behaviors of GD RBCs in response to morphological alterations and membrane stiffening. For detailed formulas and parameters to maintain RBC morphology and mechanical properties, please refer to the Supporting Materials.

**Table 1.** Model Parameters Extracted from Available Measured Data for RBCs in Normal and GD Conditions.

|  | CTR-RBC | GD-RBC1 | GD-RBC2 | GD-RBC3 |
| --- | --- | --- | --- | --- |
| Shear modulus, $G$ ($\mu N/m$) | 4.3 | 38.7 | 86.0 | 17.2 |
| RBC disaggregation threshold ($s^{-1}$) | 80 | 250 | 680 | 150 |

Notes: The mechanical properties of CTR-RBC are within the range of previous measurements [38–41]. The membrane properties for GD-RBCs are based on the experimental measurements by Franco et al. [8]. CTR-RBC represents RBCs from healthy controls, while GD-RBC1, GD-RBC2, and GD-RBC3 represent RBCs from GD patients with moderate, severe, and mild conditions, respectively,

### Cell-Cell aggregation models

In DPD method, particles interact through a series of pairwise interactions that represent the chemical and physical forces acting between different components. Details of the DPD method and the parameters adopted in this work are shown in the supplementary material.

To investigate the effect of GD RBC aggregation on blood viscosity, we applied an attractive force between particles belonging to different RBCs. These interactions are approximated with the Morse potential, defined as

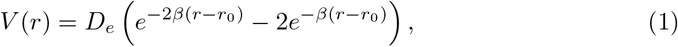

where *r* denotes the distance between two particles, *D*_*e*_ is the depth of the potential well, *β* denotes the interaction range, and *r*_0_ represents the zero-force distance. The Morse potential was applied to a specific kind of vertices of each RBC, which are called “interactive vertices” (i.e., the blue vertices in Fig 1Bi and Fig 1Bii, characterizing three different severities levels as severe, moderate and mild, as studied in [33]). To construct RBC models that reflect varying levels of cell-cell adhesion strength, we adopted the hypothesis that RBC aggregation primarily resulted from the “cross-bridging” effect, wherein each end of a sphingolipid molecule binds to two different RBCs [33, 36]. In Fig 1Bii, the presence of more blue vertices corresponds to a stronger RBC disaggregation threshold, which is commonly observed in blood with higher glucocerebroside accumulation levels. The severe condition corresponds to the highest GD RBC disaggregation threshold of 680 s^−1^ in the experiment; the moderate condition corresponds to a mean threshold of 250 s^−1^; and the mild condition corresponds to a lowest threshold of 150 s^−1^. According to the “cross-bridging” hypothesis, the strength of the pairwise connections (bridges) between “interactive vertices” is considered fixed. Consequently, a greater number of established bridges between two cells results in stronger adhesion. The adhesion strength of each bridge is mainly correlated with the parameter *D*_0_ in Eq. 1. In addition, to prevent RBC membranes from overlapping, we applied a repulsive term of Lennard-Jones potential to all membrane vertices; this potential is given by

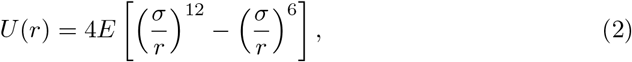

where *E* and *σ* are scaling constants for energy and distance, respectively, and these interactions vanish for *r* > 2^1*/*6^*σ*. Detailed model parameters are provided in the Supporting Materials and Methods section.

**Fig 1.**
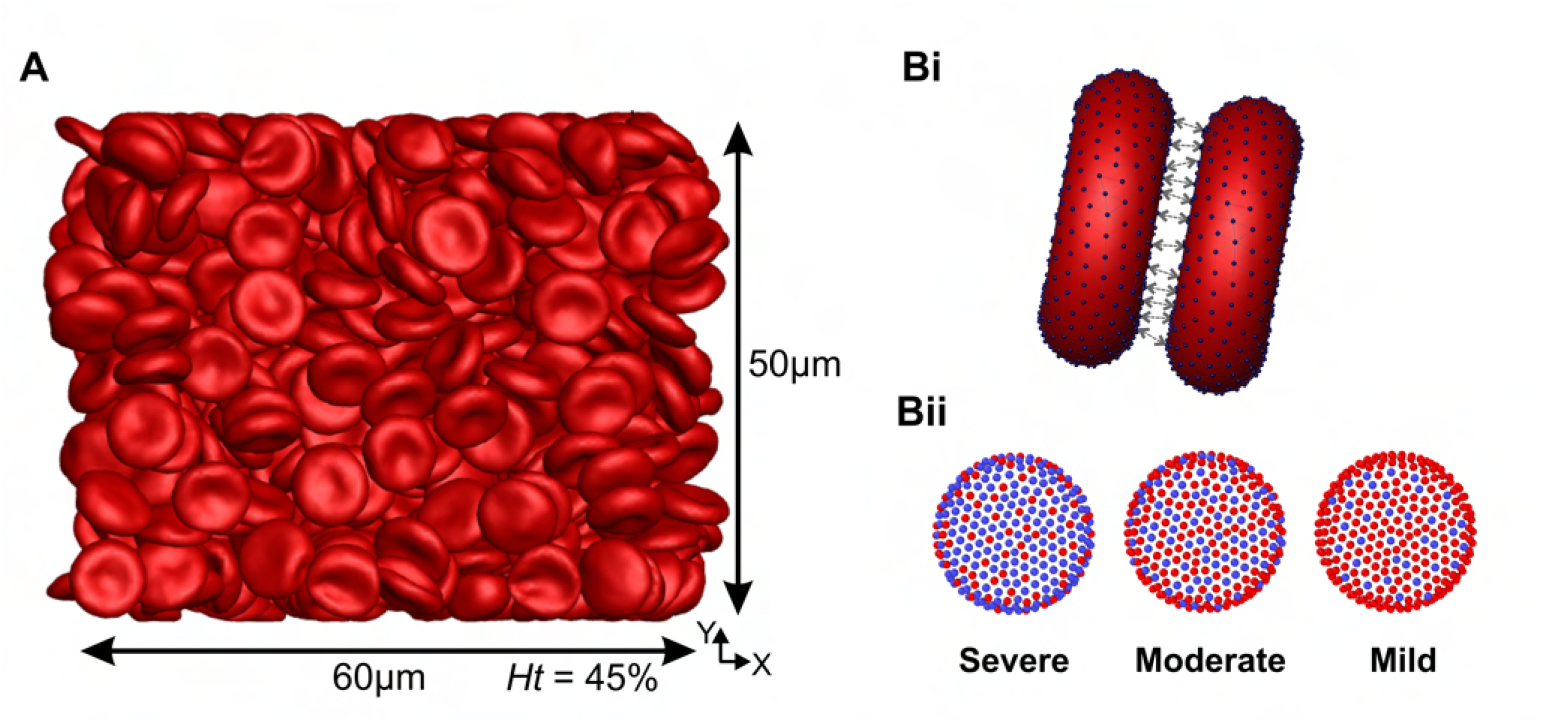
Overview of simulation setup and Cell-Cell interaction models. A) The front view of a typical simulation of blood viscosity at *Ht* = 45% (corresponding to 698 RBCs) and the size of the simulation RBC populations (60*µ*m *×* 50*µ*m *×* 50*µ*m) employed in this study. Bi) Vertex-vertex interactions for the aggregation model. Bii) Representation of three different disaggregation thresholds through the Morse potential between constituents of different RBCs (blue vertices indicate the attractive particles of RBCs). Here, the presence of more blue vertices corresponds to a stronger RBC disaggregation threshold, which is commonly observed in blood with higher glucocerebroside accumulation levels. The severe condition corresponds to the highest RBC disaggregation threshold of 680 s^−1^ in the experiment; the moderate condition corresponds to a threshold of 250 s^−1^; and the mild condition corresponds to a threshold of 150 s^−1^. Detailed model parameters are provided in the Supporting Materials and Methods section.

### Computational rheometry

In a DPD simulation, each particle represents the center of mass for a small group of atoms or molecules, with its position and momentum updated in a continuous phase but at discrete time intervals. Particles *i* and *j*, located at positions **r**_*i*_ and **r**_*j*_ respectively, occur through three types of pairwise forces: conservative, dissipative, and random forces. The viscosity is determined by dividing the calculated shear stress by the shear rate. Having individual particle velocities and each of the pairwise interactions between particles, the stress tensor, *S*, can be directly calculated through the Irving-Kirkwood formalism [37] as follows:

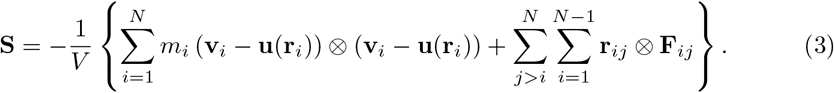

Here, *V* is the volume of the entire fluid and *u*(**r**_*i*_) is the streaming velocity imposed by the shearing flow. The ⊗ is the dyadic product of the two vectors. The shear viscosity of the DPD fluid is subsequently defined as the shear component of the stress tensor divided by the imposed deformation rate by the wall velocities

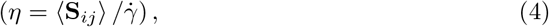

see [26]. The volume fraction of RBCs in the suspension, namely the hematocrit (*Ht*), can be calculated using the volume *V* ^*′*^ and the number of RBCs *N* in a system with the volume *V* as follows: 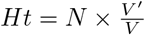. The hematocrit was set to 25 % to 45 %, matching the hematocrit observed in healthy subjects and GD patients [8]. We performed the simulations over long timescales, and the shear stress was then averaged over the entire steady-state flow. Plasma viscosity was excluded as relevant protein levels were normal [8].

### Simulation setup

In the viscosity simulations, illustrated in Fig 1A, we modeled a rectangular box with dimensions of 60 *µ*m in the X-direction, 60 *µ*m in the Y-direction, and 50 *µ*m in the Z-direction (perpendicular to the XY plane) and bounded by two walls of width 5 *µ*m. The simulation box considered in the current study (Fig 1A) has been shown to be sufficiently large for accurate rheological characterization [27]. The shear flow was imposed by the bounding walls moving in opposite directions at a constant velocity. The boundary conditions were periodic in the velocity (*X*) and the vorticity (*Z*) directions, and no-slip boundary condition was applied near the walls. Each individual RBC was composed of 500 DPD particles, with the number of RBCs modeled ranging from 155 to 698 to reproduce hematocrits of *Ht* = −10 45%. Accordingly, 600, 000 particles were modeled to represent the plasma, resulting in a total number of particles ranging from 677, 500 to 949, 000.

All simulations were computed by using the extended version of code developed based on LAMMPS. Each simulation takes about 2,000,000 time steps to 4,000,000 time steps. A typical simulation requires 2400 CPU core hours to 4800 CPU core hours by using the computational resources (Intel Xeon E5-2670 2.6 GHz 24-core processors) at the Center for Computation and Visualization at Brown University.

### Experiment setup

We used data from different experiments previously done in Franco et al. [8], approved by the local ethics committee in accordance with the principles of declaration of Helsinki. These experimental data form the basis of this study: adhesion assay on laminin at 1 dyne/cm^2^; RBC aggregation analysis by syllectometry; and blood viscosity measured by cone/plate viscometer.

For GD RBC deformation during the adhesion assay, the experiment was performed using a plate flow chamber and an ibiTreat-coated or uncoated microslide I^0.2^ Luer (Ibidi) [8]. Ten *µ*g of laminin *α*5 chain was immobilized on an uncoated microslide and RBCs, resuspended at 0.5% hematocrit (Hct), were perfused through the microslide under a 0.2 dyn/cm^2^ flow for 5 minutes at 37°C. After cells were washed out, RBCs were fixed at 1 dyn/cm^2^, and RBC morphology and Lu/BCAM pattern were visualized by immunofluorescence using a specific antibody and confocal microscopy [8]. We qualified the shear modulus for GD RBCs by determining the X-axis length of CTR and GD RBCs under a shear stress of 1 dyn/cm^2^ from the confocal images of adherent CTR and GD RBCs on laminin *α*5. The X-axis length measurements were performed on several images obtained from adhesion assay experiments conducted at a shear stress of 1 dyn/cm^2^, using CTR (*n* = 10) or GD (*n* = 11) RBCs. The X-axis corresponds to the flow direction. The mean X-axis length of CTR RBCs is 11.0 *µ*m (*n* = 10), while the mean X-axis length of GD RBCs is 8.9 *µ*m (*n* = 11) (Fig 2B).

**Fig 2.**
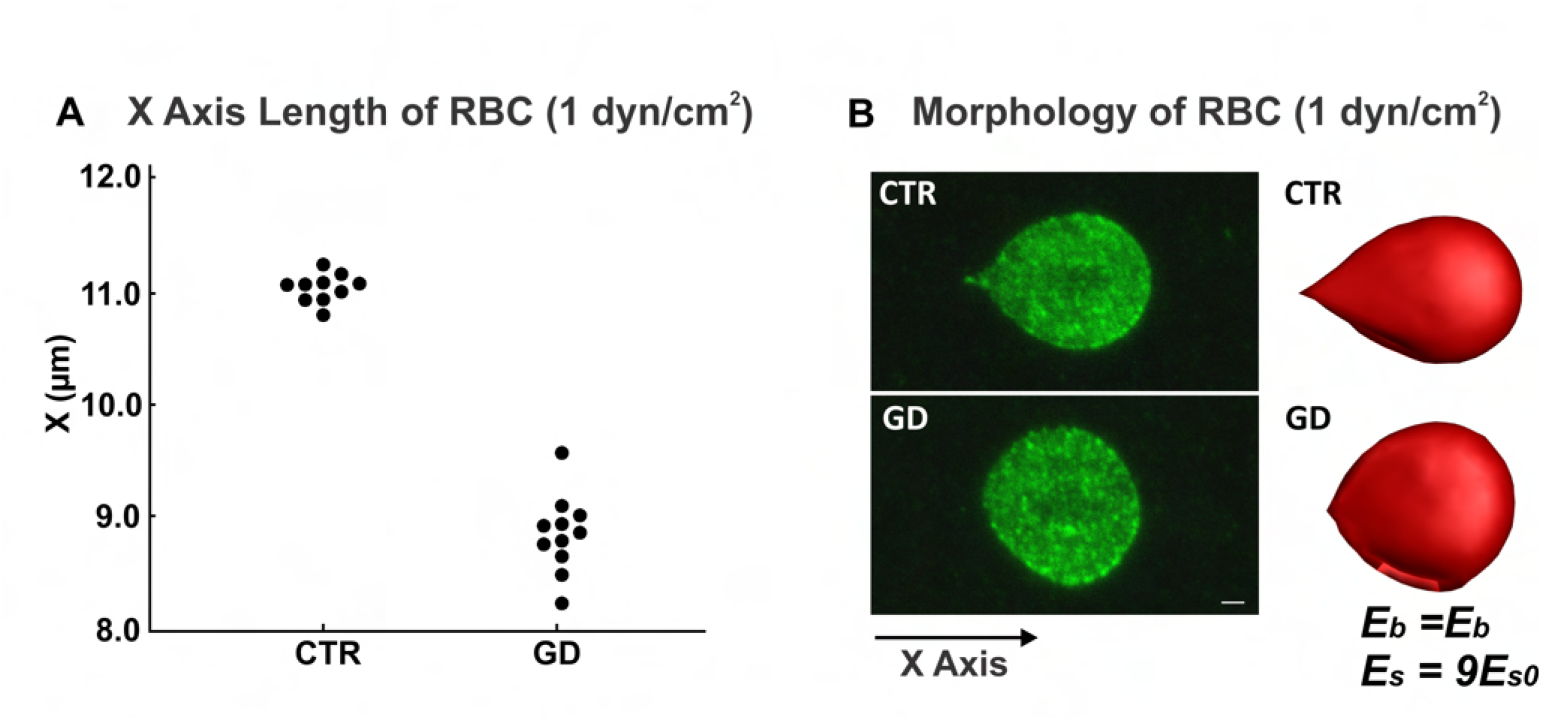
Validation of shear modulus of GD RBC with adhesion assay. A) X-axis length of CTR and GD RBCs from experiments at a shear stress of 1 dyn/cm^2^. The X-axis length measurements were performed on several images obtained from adhesion assay experiments conducted at a shear stress of 1 dyn/cm^2^, using CTR (*n* = 10) or GD (*n* = 11) RBCs. B) Morphology of CTR and GD RBC under shear flow. The left two figures are confocal images of adherent CTR and GD RBCs to laminin *α*5 at a shear stress of 1 dyn/cm^2^. The right two figures are simulation images for CTR and GD RBCs at different shear modulus conditions at a shear stress of 1 dyn/cm^2^. Bar scale represents 1 *µ*m. The left figure of the panel (B) was originally published in Blood, Franco, et al. 2013 Jan [8], © the American Society of Hematology.

The RBC aggregation experiment was conducted at 37°C by syllectometry (i.e., laser backscatter versus time, using the Laser Optical Rotational Red Cell Analyzer; LORRCA, Mechatronics, The Netherland) after adjustment of the hematocrit to 40%. The RBC disaggregation threshold (*γ*), that is, the minimal shear rate needed to prevent RBC aggregation or to breakdown existing RBC aggregates, was determined using a reiteration procedure [8].

Blood viscosity experiment was setup at native Hct, at 37°C, and at various shear rates using a cone/plate viscometer (Brookfield DVIII+ with CPE40 spindle, Brookfield Engineering Labs, USA). All hemorheological measurements were carried out within 4 hours of venipuncture to avoid blood rheological alterations and after complete oxygenation of the blood. We followed the guidelines for international standardization of blood rheology techniques/measurements and interpretation [42].

## Results

In this section, we first validated the simulation parameters against experimental data. Then, we simulated suspensions of RBCs that replicate the experimental conditions of GD erythrocyte suspensions as conducted by Franco et al. [8]. Last, we investigated the effects of hematocrit level, RBC deformability, and RBC disaggregation threshold on blood viscosity by DPD simulation.

### Clinical and hemorheological profile of GD patients with bone disease

This study focused on acute or subacute manifestations of GD such as bone infarcts and AVN. We conducted an analysis on 22 untreated GD patients, among whom 7 exhibited various forms of bone disease, including bone infarcts and vertebral body collapse, indicating a prevalence of bone disease in 31.8% of GD patients. Among these 22 patients, hemorheology experiments were performed on 9 patients [8]. Table 2 provided a detailed overview of the clinical and biological parameters of 3 GD patients who have been diagnosed with bone disease. Each RBC parameter from these 3 patients was compared with mean values calculated from 11 healthy volunteers (M-CTR) and from 9 GD patients (M-GAU) (Table 2). Notably, both Patient 1 and Patient 3 demonstrated substantially lower elongation indices (calculated as

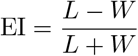

where *L* is the cell’s length along the flow axis and *W* its width perpendicular to the flow) compared to the M-CTR and M-GAU across shear stress levels ranging from 0.3 Pa to 3.0 Pa. Patient 2 exhibited lower elongation index values compared to the M-GAU values at shear stress level ranging from 0.53 to 0.3 Pa, indicating a significant decrease in RBC deformability. Furthermore, all three patients display higher RBC disaggregation thresholds and blood viscosities compared to the M-CTR, with only Patient 1 exceeding the M-GAU threshold. These data were insufficient to conduct robust statistical analysis regarding potential links between hemorheological parameters and acute/subacute bone disease in GD. However, the markedly decreased elongation indices, elevated RBC disaggregation thresholds, and increased blood viscosity suggest underlying biophysical mechanisms in these complications encouraging a deeper exploration of biophysics and rheology of blood and RBCs in GD.

**Table 2.** Clinical and biological parameters of the studied patients with hemorheology experiment.

| Patient No. | 1 | 2 | 3 | M-CTR | M-GAU |
| --- | --- | --- | --- | --- | --- |
| Clinical skeletal manifestations | - | - | bone pains | - | - |
| Radiological (MRI) skeletal manifestations | VBC | BI | BI | - | - |
| Hb (g/dl) | 13.9 | 12.5 | 12.0 | - | 12.6 |
| EI at 0.3 Pa | 0.004 | 0.043 | 0.002 | 0.055 | 0.021 |
| EI at 0.53 Pa | 0.055 | 0.055 | 0.053 | 0.084 | 0.062 |
| EI at 0.95 Pa | 0.113 | 0.114 | 0.123 | 0.165 | 0.131 |
| EI at 1.69 Pa | 0.202 | 0.216 | 0.226 | 0.262 | 0.229 |
| EI at 3.0 Pa | 0.303 | 0.317 | 0.331 | 0.356 | 0.329 |
| RBC disaggregation threshold ($s^{-1}$) | 800 | 190 | 215 | 111 | 245 |
| Blood viscosity at $22.5 s^{-1}$ | 11.85 | 7.46 | 8.17 | 8.03 | 9.01 |
| Blood viscosity at @ $45 s^{-1}$ | 9.6 | 6.39 | 6.9 | 6.62 | 7.43 |
| Blood viscosity at @ $90 s^{-1}$ | 7.89 | 5.39 | 5.93 | 5.74 | 6.32 |
| Blood viscosity at @ $225 s^{-1}$ | 6.26 | 4.9 | 4.86 | 4.98 | 5.40 |

Notes: M-CTR: Mean value of 11 healthy volunteers; M-GAU: Mean value of 9 GD patients; BI: patients whose skeletal MRI revealed at least 1 bone infarct, either painful or not in the disease history; VBC: vertebral body collapse; Hb: hemoglobin; EI: Elongation Index; rpm: revolutions per minute of viscometer.

### Calibration of the GD RBC deformability during adhesion assay

Membrane deformation of GD RBCs during flow adhesion experiment was visualized by confocal microscopy. At 1 dyn/cm^2^, CTR RBCs were elongated and underwent a deformation of their membrane, with RBCs exhibiting a typical racket shape described by Franco et al. [8]. In contrast, GD RBCs did not change shape, suggesting reduced membrane deformability (left column of Fig 2B).

In analogy with the experimental setups [8], simulations were conducted where a single RBC was initially attached to a substrate, and a pressure gradient was imposed on the liquid, inducing shear flow on the RBC. The direction of the shear flow followed the X-axis, from left to right, as shown in Fig 2B. Subsequently, a stepwise increasing pressure gradient was applied to the liquid until the X-axis length of the CTR RBC in the simulation matched that of the CTR RBC in the experiment, i.e., 11.6 *µ*m. At this point, the simulated RBC and the experimental RBC exhibited similar morphology (top column of Fig 2B). We determined that the pressure gradient parameter in this simulation corresponded to a shear stress of 1 dyn/cm^2^. After establishing this shear stress, we maintained it at 1 dyn/cm^2^ while varying the shear modulus of the RBC in simulations from 1*Es*_0_ to 25*Es*_0_. After completing the simulations for each shear modulus of the GD RBC, we compared the X-axis length of each RBC under shear stress to the X-axis length of the GD RBC in the experiment (Fig S1). We found that the RBC with a shear modulus of 9*Es*_0_ closely matched the experimental GD RBC in terms of X-axis length and morphology. The morphology of the RBC in the simulation closely resembled that of the RBC observed in the experiment, indicating that the simulation parameters accurately reflected the experimental conditions (bottom row of Fig 2B).

### Calibration of the aggregation model based on syllectometry

In analogy with experimental setups [8], simulations were conducted with two RBCs initially in contact one with another forming a doublet. After sufficient contact time and the cell-cell interaction reaching an equilibrium, we imposed the bounding walls moving in opposite directions at a constant velocity. The shear flow was induced by the moving of bounding walls. Subsequently, we imposed a stepwise increasing shear rate on the bounding walls until the doublet broke up, i.e., these two RBCs were fully detached from each other. In the simulation, cell-cell aggregation between the two stacked RBCs occurred after they approached each other. The setup involved two parallel surfaces moving in opposite directions to generate shear flow along the horizontal direction. As a result, the two aggregated RBCs were sheared apart by sliding (Fig 3). By using this method, we could calibrate the pairwise interaction strength in cell-cell aggregation models with the RBC disaggregation threshold observed in the experiment.

**Fig 3.**
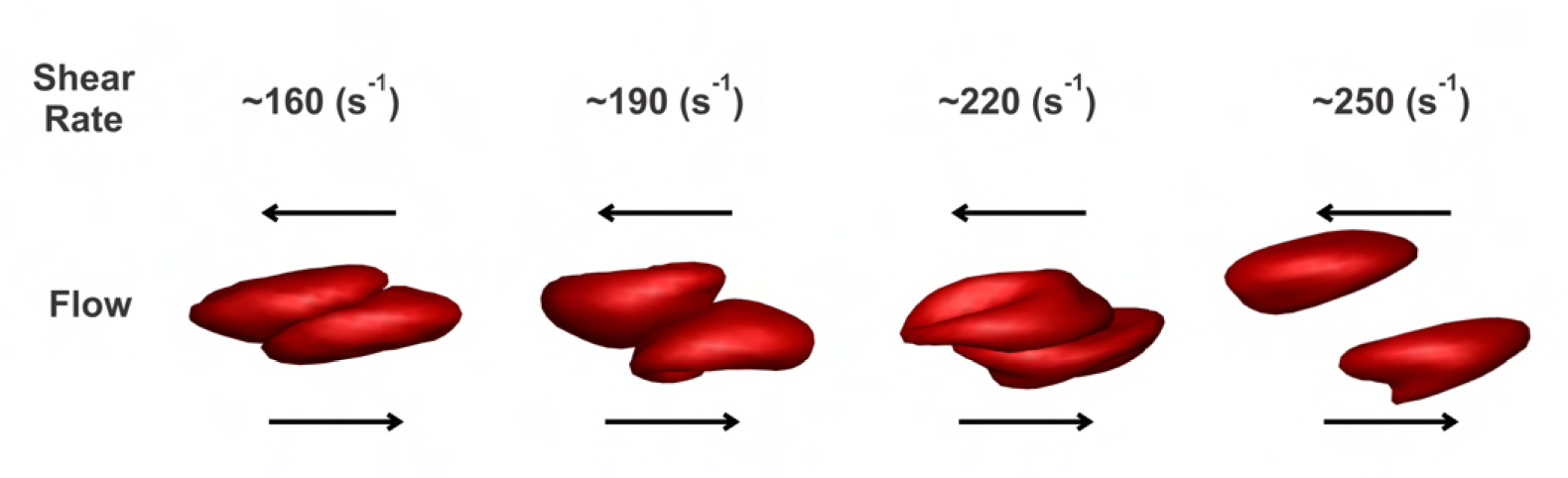
Sequential snapshots of the disaggregation dynamics of GD RBCs (GD-RBC1) for a range of varied flow shear rates.

To simulate the cell-cell detachment of GD RBC doublets under different shear rates, we also carried out the simulations on the breakup of GD RBC doublets and recorded the minimal shear rate that allowed the full separation of these two RBCs in doublets (Fig 3). Generally, the cell-cell disaggregation threshold increased as the interactive vertices ratio (aggregate severity) increased in both viscometer experiments and simulations. Also, larger *D*_*e*_ introduced the condition that a higher shear rate was required to break up the doublets, i.e., The cases with *D*_*e*_ = 4.24 × 10^−25^ J required higher detaching shear rates compared with those with *D*_*e*_ = 8.48 × 10^−26^ J. The increasing RBC disaggregation threshold required to fully detach RBC doublets in the simulation closely resembled the results from cone/plate viscometer experiments, indicating that simulation parameters accurately captured the disaggregation dynamics observed experimentally (Fig 3).

### Modeling of the rheology of GD RBC suspensions

Franco et al. demonstrated increased bulk viscosity in GD RBC suspensions under physiological shear rates (22.5–225 s^−1^) using a cone/plate viscometer, [8]. For the *in silico* viscosity measurement, we used our model, taking into account non-Newtonian behavior such as shear thinning and the Fahraeus–Lindqvist effect [43].

To evaluate the accuracy of our *in silico* viscosity measurements, we compared our values generated with viscosities measured by Franco et al. [8]. Fig 4 shows the steady shear blood viscosity as a function of shear rate, which shows an excellent agreement between the simulation results and the experimental measurements for normal and GD RBC suspensions [8].

**Fig 4.**
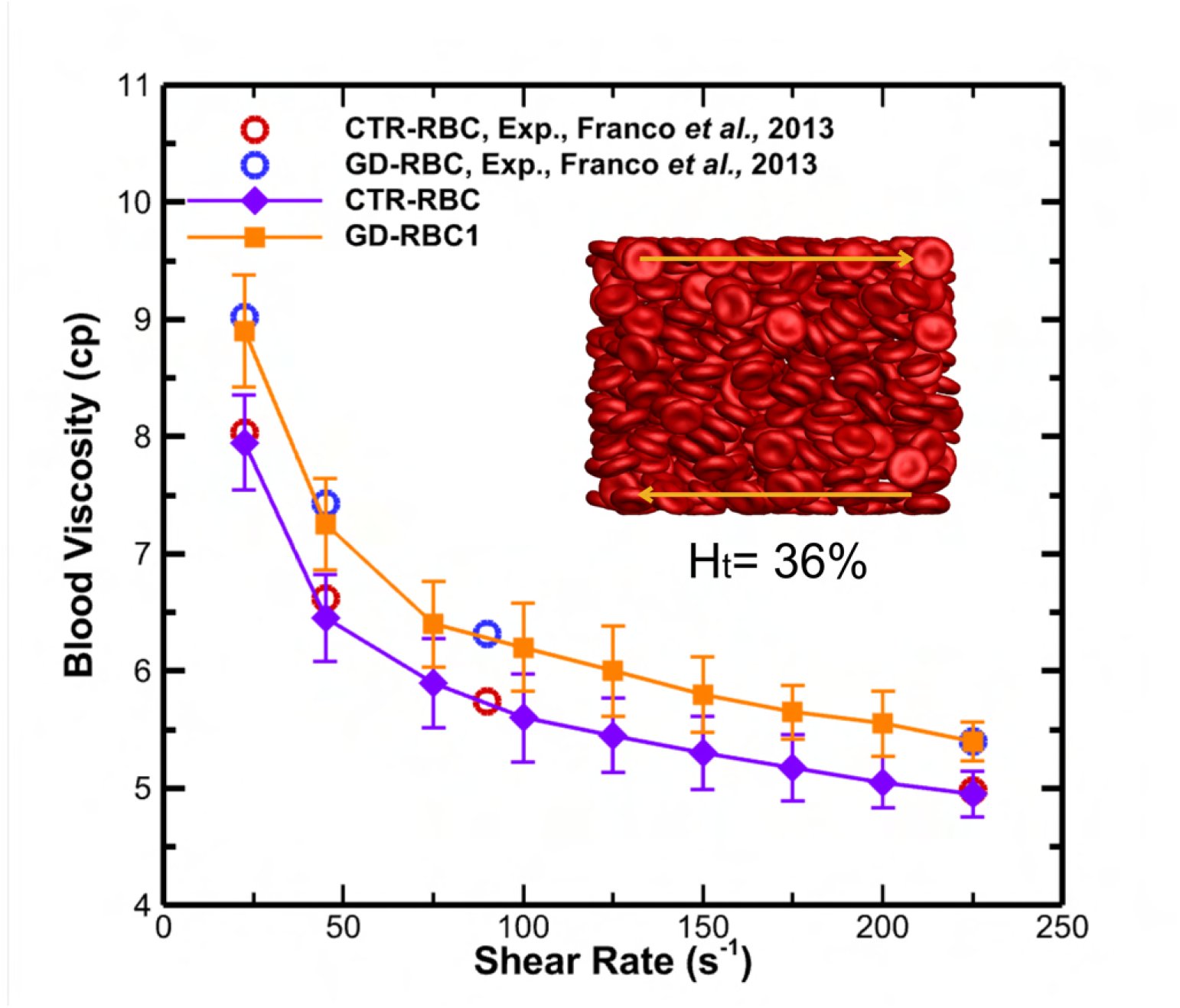
Blood relative viscosity for GD evaluated in viscometric flows under physiologically relevant shear rates ranging from 22.5 to 225 s^−1^. Experimental viscosity measurements for GD and CTR RBC suspension by Franco et al. [8]. The viscosity measurements were based on a hematocrit of ∼36% for GD RBCs and ∼40% for CTR RBCs. Open symbols represent experimental results, whereas solid symbols represent simulation results.

We then extended our analysis to examine the relative blood viscosity in GD under shear flows with larger shear rates, ranging from 1 to 1000 s^−1^. This four-orders-of-magnitude range covers most of the shear levels present in the micro- and macrocirculation, from low/moderate rates in veins (around 1 s^−1^) and arteries (around 80 ™ 250 s^−1^) to high rates (up to 1000 s^−1^) in capillaries and arterioles [44–46]. In Fig 5, viscosity curves were plotted for normal RBC (CTR-RBC) and GD-RBC1, GD-RBC2, and GD-RBC3, which represent RBCs from GD patients with moderate, severe, and mild conditions, respectively, as shown in Table 1. These curves revealed three distinct regions that we have categorized as separate domains.Under physiologically high shear rates (*>* ∼100*s*^−1^), the elevated blood viscosity of GD patients seems to be representative of the blood rheological anomalies of various vascular diseases such as atherosclerosis [47], and malaria-infected blood [48]. Under physiologically lower shear rates (⩽ ∼8*s*^−1^), the blood viscosity of GD patients could also be representative of these diseases, particularly in diabetic mellitus because of the RBC hyperaggregation [49].

**Fig 5.**
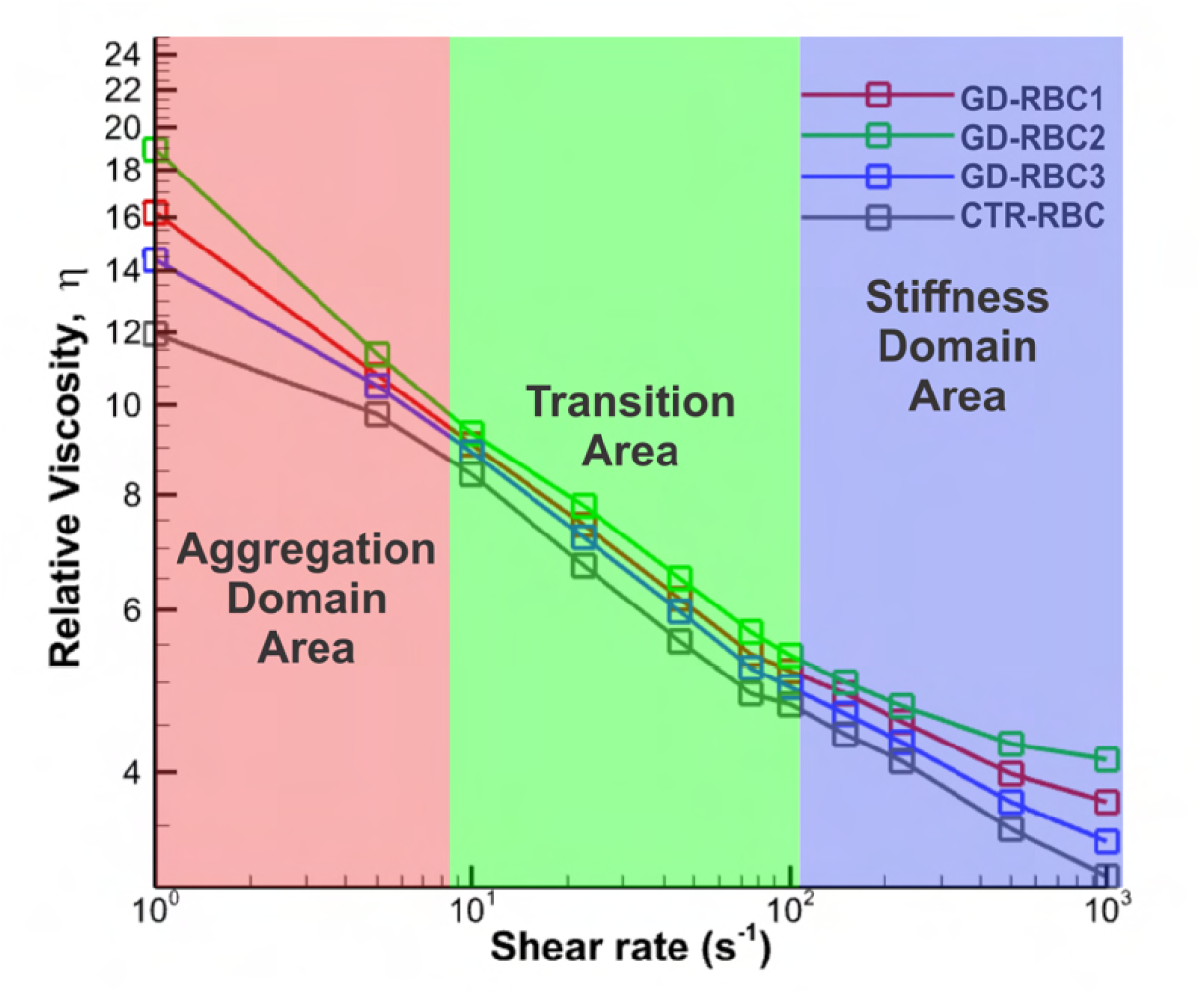
Extension of blood relative viscosity for GD in shear flows under large shear rates ranging from 1 to 1000 s^−1^. There are three distinct domains for blood viscosity based on the shear rate: the aggregation domain, ranging up to approximately 8 s^−1^; the transition area, spanning from around 8 to 100 s^−1^; and the stiffness domain, extending from approximately 100 s^−1^.

To investigate whether different underlying mechanisms dominate across varying flow domains in GD blood, similar to other blood rheological anomalies, we defined three distinct domains of GD blood viscosity based on shear rate. Then, we investigated whether different underlying mechanisms dominate across varying flow domains in blood from GD patients.

#### 0.0.1 Impact of the RBC hyperaggregation in GD patients on blood viscosity

Here, we studied the role of the RBC disaggregation threshold on blood viscosity by increasing the number of RBC constituents that are attractive to one another which will induce inter RBCs attachments. Fig 6A shows the schematic representation of different RBC disaggregation thresholds through the Morse potential between constituents of different RBCs. This methodology accurately represents the increased effective interactions without prohibitive computational costs, which is consistent with the results of Deng et al. [50]. Fig 6B shows the relative viscosity of blood for different hematocrit levels (Ht = 25%, 36%, 45%) over a range of shear rates (1 to 1000 s^−1^) for different RBC disaggregation thresholds. The shear rates and hematocrits examined in our study were consistent with the physiological ranges observed in the circulatory system of GD patients [8, 51]. Here, we added an intermediate threshold of 350 s^−1^ to reduce the large gap between 250 s^−1^ and 680 s^−1^. The relative viscosity plotted revealed a stark increase of viscosity at low shear rates for both hematocrit levels and RBC disaggregation thresholds studied. The observed variation in viscosity between the different colored curves is due to the different hematocrit levels tested, which shows that a high hematocrit level would contribute to high blood viscosity (Fig 6B). The discrepancy in the same color is caused by the RBC disaggregation threshold, which is much more significant at low shear rates. In contrast, virtually no change is observed at larger shear rates. Thus, the first domain of the curve is dependent on the aggregation properties of the GD RBCs.

**Fig 6.**
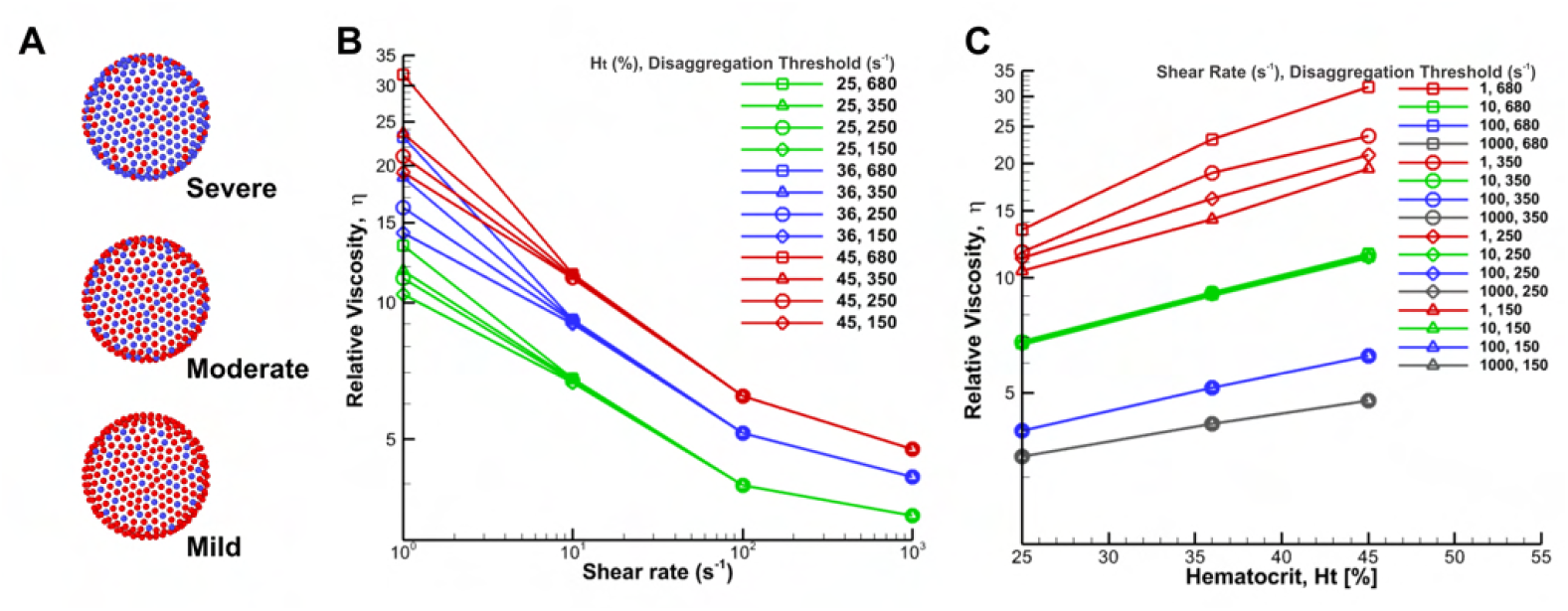
**Schematic diagram of different RBC disaggregation thresholds (A) and corresponding simulation results (B and C) of the relative viscosity of blood for different RBC disaggregation thresholds and hematocrit levels with shear modulus of** 9*E*_*s*0_. A) Representation of three different glucocerebroside accumulation through the Morse potential between constituents of different RBCs. Here, the presence of more blue vertices corresponds to a stronger RBC disaggregation threshold. The severe condition corresponds to the highest RBC disaggregation threshold of 680 s^−1^ in the experiment; the moderate condition corresponds to a threshold of 250 s^−1^; and the mild condition corresponds to a threshold of 150 s^−1^, as defined in Table 1.; B) The steady shear relative viscosity versus imposed shear rate for three different hematocrit levels and four different RBC disaggregation thresholds at each hematocrit. The viscosities are normalized by the plasma viscosity; the effect of RBC disaggregation thresholds disappears at high shear rates when it causes no stable cell-cell adhesion; C) Relative viscosity against hematocrit for different shear rates and four different RBC disaggregation thresholds of 150, 250, 350 and 680 s^−1^. In the simulation, we added an intermediate threshold of 350 s^−1^ to reduce the large gap between 250 s^−1^ and 680 s^−1^.

Additionally, the increase in viscosity at low flow rates was significantly more pronounced at higher hematocrit values. This is clearly illustrated by the red lines in Fig 6C, where the relative viscosity is plotted against hematocrit for various shear rates and RBC disaggregation thresholds. In contrast, when the relative viscosity is plotted at high flow rates, the resulting lines overlap, indicating that the disaggregation threshold has no significant influence on viscosity at higher shear rates, regardless of the hematocrit level (blue and grey lines in Fig 6C).

#### 0.0.2 Impact of the cell deformability on blood viscosity in GD

Here, we directly manipulated the deformability of RBCs in isolation by adjusting the shear modulus that maintains RBC mechanical integrity. The viscosity is reported for three distinct cases of RBC deformability/shear moduli in Fig 7. Fig 7A depicts the representation of three different levels of cell stiffness through different shear modulus conditions at a shear stress of 1 dyn/cm^2^. Fig 7B and Fig 7C illustrate an overall increase in viscosity across different hematocrits and shear rates, with this augmentation notably more pronounced under physiologically higher shear rates (100–1000 s^−1^, see red lines in Fig 7B and grey lines in Fig 7C). We also noted that blood viscosity in GD, as predicted by the model, rises by approximately 1.4-fold compared with normal values when hematocrit increases from 25% to 45% (Fig 7B and Fig 7C). Then, the third domain of the curve is dependent on the deformability of the RBCs.

**Fig 7.**
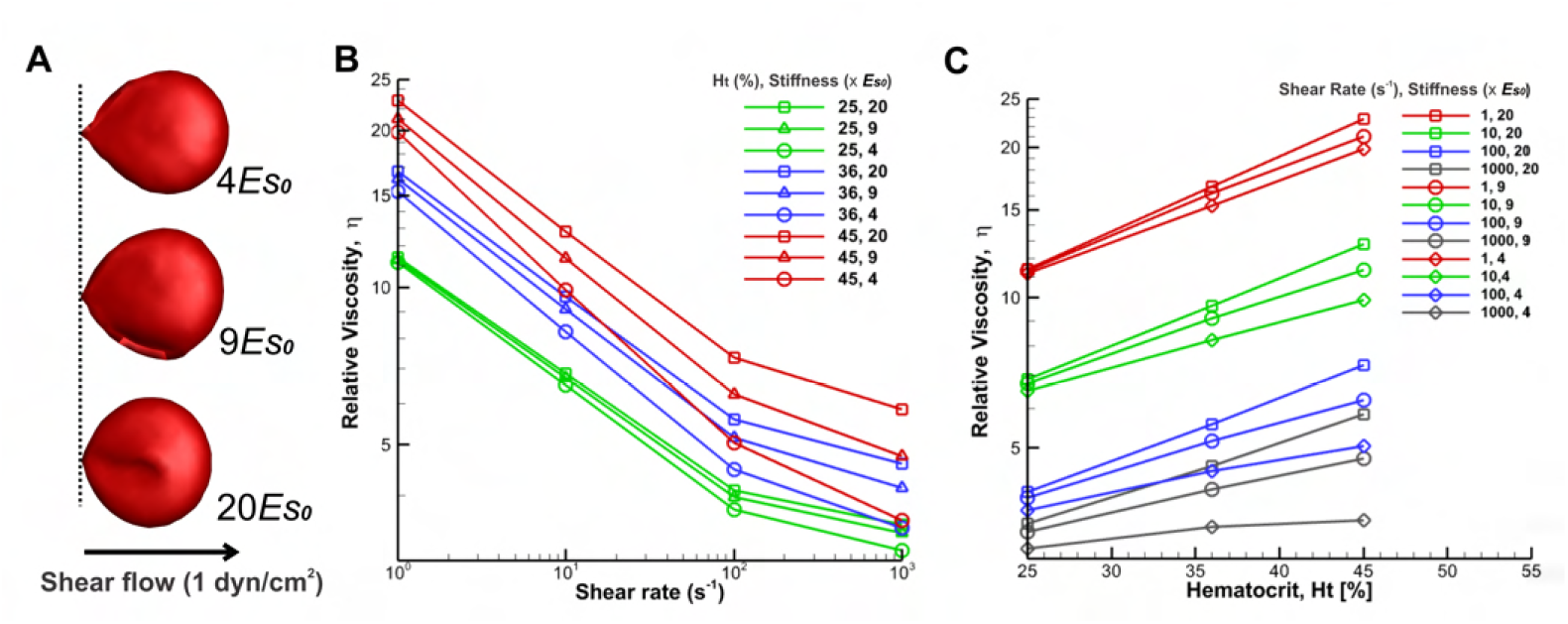
**Schematic diagram of different RBC stiffness (A) and corresponding simulation results (B and C) of the relative viscosity of blood for different RBC stiffness and hematocrit levels with disaggregation threshold of** 250*s*^−1^. A) Representation of three different levels of cell stiffness through different shear modulus conditions at a shear stress of 1 dyn/cm^2^. B) The steady shear relative viscosity versus imposed shear rate for three different hematocrit levels and three different cell stiffness at each hematocrit. The viscosities are normalized by the plasma viscosity; the effect of cell stiffness increases at high shear rates, and the deformation of cells becomes increasingly more dominant C) Relative viscosity against hematocrit for different shear rates and three different RBC stiffness of 20, 9, and 4 *E*_*s*0_.

At very low shear rates, blood rheology and flow dynamics are predominantly influenced by fluid flow and stress fluctuations between particles, explainable through suspension mechanics. Conversely, at higher shear rates, cell deformation becomes increasingly dominant. Thus, blood containing RBC with decreased deformability exhibits heightened viscosities at larger shear rates compared to blood containing normally deformable RBC. This effect is stronger at higher hematocrit levels.

#### 0.0.3 Impact of the hematocrit levels on blood viscosity in GD

In this section, we investigated the impact of hematocrit levels on blood viscosity, in combination with the deformability and aggregation properties of GD RBCs. Thus, we changed the hematocrit from 25% to 45% and explored the non-Newtonian viscosity over a range of shear rates in GD. Fig 6 and Fig 7 present the steady shear relative blood viscosity as a function of shear rate and hematocrit. As generally observed in physiology and disease, at all shear rates blood viscosity increased dramatically with rising hematocrit, as shown in Fig 6C and Fig 7C; however, this increase also depended on the shear rate at which the viscosity was measured. For instance, in Fig 7C, increasing the hematocrit from 25 to 45% at the shear rate of 1 s^-1^ resulted in a 1.8-fold higher value of blood viscosity, whereas at a shear rate of 1000 s^-1^, it only increased 1.4-fold the value of blood viscosity. Thus, the effect of hematocrit on blood viscosity is greater at low shear rates.

In GD patients, hematocrit levels are often below the normal range, so we expected decreased blood viscosity. The average hematocrit level in blood samples from 11 untreated GD patients was 36.42% ± 3% (n = 11) [8]. This anemia can be explained by the dyserythropoiesis observed in GD and by the splenic retention of rigid GD RBCs, which results in a drop in hematocrit. However, treated patients may have a hematocrit level of 40% - 45%. At low shear rates, this increase in hematocrit to 45% may result in blood viscosity roughly doubling compared to that of healthy individuals. Similarly, at high shear rates, the viscosity may rise to approximately 1.8 times that of healthy individuals (Fig 6B). The hematocrit levels in Fig 6 and Fig 7 (25% - 45%) are within the physiological range for GD. While hematocrit significantly impacts blood viscosity, other factors also contribute to the overall viscosity profile.

## DISCUSSION AND SUMMARY

Bone complications in GD are of multiple types and mechanisms, including bone fragility marked by osteopenia/osteoporosis, osseous deformities, and cortical thinning in long bones [9, 10, 52, 53]. Additionally, acute or subacute occurrences such as bone infarcts and AVN of the head of long bones or vertebral bodies contribute to its complexity. Bone infarcts and AVN occur when bones do not get enough blood due to decreased intraosseous blood flow, causing the tissue to deteriorate and die, resulting in tiny breaks in the bone and eventual bone collapse [10, 11]. Although reminiscent of a similar complication in sickle cell disease, where microcirculation impairment is central [12], the mechanisms that contribute to bone involvement in GD are not clearly established. Several studies have shown that bone pathology in GD is attributed to infiltration of the bone marrow and trabecular bone by Gaucher cells [9, 13]. On the other hand, it has been shown in different pathologies that blood viscosity abnormalities can induce ischemic processes [54, 55]. Detailed analysis of rheological data from 3 Gaucher patients with bone damage suggests that the RBC aggregation properties, as well as their deformability and blood viscosity, could play a role in the onset of bone damage. Furthermore, a large study from the International Gaucher Disease Registry (ICGG) indicates that patients with GD who have undergone surgical removal of the spleen (splenectomy) have a higher incidence of AVN [18]. Given that the spleen is the organ that removes biophysically altered RBCs from circulation, these findings strongly suggest a potential link between AVN occurrence and altered RBC biophysics and blood viscosity [56, 57].

Based on the rheological alterations of RBCs in GD, we hypothesized that the pathogenesis of bone infarcts and AVN in GD may involve rigid erythrocytes and elevated RBC aggregation, leading to increased blood viscosity and microcirculatory obstruction [58–62]. It would therefore seem interesting to study more precisely the impact of abnormal Gaucher RBC properties, such as aggregation and deformability, on blood viscosity. Due to the limitations of experimental *in vitro* measurements mentioned earlier, we chose to use computational methods, as advances in computational power over the past two decades have stimulated the development and applications of multiscale biophysical models [23, 63–68]. Based on previous results obtained *ex vivo* [8], in this study, we investigated the biomechanics and rheology of blood and RBCs in GD under various flow conditions via *in silico* models.

In our study, we identified three factors that affect blood viscosity in GD: RBC deformability, RBC disaggregation threshold, and hematocrit level. Firstly, experimental findings have shown that hardened RBCs are present in GD patients [8], and these cells exhibit increased resistance to flow compared to healthy ones, leading to elevated blood viscosity [19]. Reduced RBC deformability is also observed in conditions like diabetes mellitus, where shear moduli are twice those of healthy cells, contributing to higher viscosity across various shear rates [69]. Similarly, malaria-infected RBCs lose deformability, further increasing viscosity [70], while in sickle cell anemia, deoxygenated sickled RBCs can become 100–2000 times more rigid, significantly raising blood viscosity [71]. Secondly, the mechanisms underlying RBC hyperaggregation in GD patients remain unclear but likely involve cellular factors [17, 72]. Glucocerebroside, a sphingolipid that accumulates in RBC membranes, may play a key role by raising the RBC disaggregation threshold and promoting rouleaux formation, where cells stack in chain-like structures [17]. Lastly, hematocrit levels are lower in GD patients compared to healthy controls [20], which highlights the importance of considering hematocrit variations when assessing splenectomy in these patients. Post-splenectomy hemorheological changes have been linked to bone complications [73–75].

To investigate the effects of RBC deformability, RBC disaggregation threshold, and hematocrit level on blood viscosity by DPD simulation, we first validated the simulation parameters against experimental data. This study is the first to investigate the Young’s modulus and aggregation parameters of GD RBCs by integrating validated simulations with confocal imaging and experimental RBC disaggregation thresholds. Our simulation parameters accurately captured the shear modulus of GD RBCs and the RBC disaggregation dynamics observed experimentally. This approach enables a comprehensive assessment of the mechanical properties of GD RBCs.

To validate the *in silico* viscosity model against experimental measurements, we simulated suspensions of RBCs that replicate the experimental conditions of GD erythrocyte suspensions as conducted by Franco et al. [8]. Our simulation shows an excellent agreement with the experimental measurements. After that, we extended our analysis to examine the relative blood viscosity in GD with larger shear rates ranging from 1 to 1000 s^−1^.

Furthermore, to explore whether different underlying mechanisms dominate across varying flow domains in GD blood, we defined three distinct domains of GD blood viscosity based on shear rate. These domains include the aggregation domain, which spanned shear rates up to approximately 8 *s*^−1^; the transition area, which covered shear rates from around 8 to 100 *s*^−1^; and the stiffness domain, which extended from approximately 100 *s*^−1^ and beyond. The stiffness domain may correspond to the regions where RBC deformability becomes a major factor. In GD, altered RBCs could lead to reduced deformability and the ability to navigate through small capillaries, worsening the microcirculatory problems and contributing to increased vascular resistance and even organ dysfunction, such as bone infarction.

Finally, we investigated the effects of RBC deformability, RBC disaggregation threshold, and hematocrit level on blood viscosity. Our study demonstrates that all three mechanisms can cause blood viscosity to rise many folds, with manifestation in different flow rate conditions and hematocrit levels. An increased RBC disaggregation threshold leads to higher blood viscosity in GD patients at low shear rates. Consequently, this phenomenon is less likely to occur in arterioles and capillaries, which experience high shear rates. In terms of RBC deformability, decreased RBC deformability can cause elevation of blood viscosity, this can be captured under large shear rates, and observed no change at low shear rates. However, this phenomenon is less common in arteries and veins, which typically experience lower shear rates.

In summary, we identified three viscosity domains based on shear rate, showing that elevated hematocrit, reduced RBC deformability, and increased RBC disaggregation threshold contribute to increased blood viscosity with manifestation in different flow rate conditions in GD. In GD patients, lowering hematocrit helps protect the circulatory system from complications associated with high blood viscosity. Treated patients may reach hematocrit levels of 40%–45%, which could result in high blood viscosity if RBC anomalies were maintained. Fortunately, treatment significantly reduces RBC abnormalities, and as splenomegaly decreases, the spleen regains functionality, aiding in the return of blood viscosity to normal [76]. However, splenectomy can negatively impact this process by increasing hematocrit and releasing hyperaggregated, rigid GD RBCs into circulation. This underscores the role of “saving the spleen” objective in maintaining optimal blood rheology in GD patients. This study enhances our understanding of the biophysics and rheology of blood and RBCs in GD, offering insights into the broader implications of these factors in pathological conditions.

Our study of blood biophysics and rheology in GD has some limitations. While RBC membrane parameters were calibrated with experimental data, assuming fixed shear modulus and RBC disaggregation thresholds may not account for variability across individuals or GD stages. Additionally, the model reflects RBC dynamics in controlled environments, which may not fully capture *in vivo* conditions where factors like immune cell interactions affect blood rheology. Further mapping these simulations to real physiological contexts could provide deeper insights into GD-related hemorheological phenomena.

In future work, a detailed study of cell alignment, the relation between EI and RBC stiffness *in silico*, different shapes at varying hematocrits and deformation rates, and mapping of such structures with respect to different biophysical factors at play will provide an invaluable insight into the groundwork of hemorheological phenomena of GD.

## Supporting information

**S1 Text. In silico biophysics and rheology of blood and red blood cells in Gaucher Disease**.

**S2 Text. Cell-cell interaction potentials and measurements of membrane stiffness**.

## Acknowledgments

We acknowledge support from the National Institutes of Health (Grant No. R01HL154150) and the Laboratory of Excellence GR-Ex (Grant ANR-11-LABX-0051). Simulations were carried out at the Center for Computation and Visualization of Brown University.

